# Midkine-a functions as a universal regulator of proliferation during epimorphic regeneration in adult zebrafish

**DOI:** 10.1101/2020.04.14.040972

**Authors:** Nicholas B Ang, Alfonso Saera-Vila, Caroline Walsh, Peter F. Hitchcock, Alon Kahana, Ryan Thummel, Mikiko Nagashima

## Abstract

Zebrafish have the ability to regenerate damaged cells and tissues by activating quiescent stem and progenitor cells or reprogramming differentiated cells into regeneration-competent precursors. Proliferation among the cells that will functionally restore injured tissues is a fundamental biological process underlying regeneration. Midkine-a is a cytokine growth factor, whose expression is strongly induced by injury in a variety of tissues across a range of vertebrate classes. Using a zebrafish Midkine-a loss of function mutant, we evaluated regeneration of caudal fin, extraocular muscle and retinal neurons to investigate the function of Midkine-a during epimorphic regeneration. In wildtype zebrafish, injury among these tissues induces robust proliferation and rapid regeneration. In Midkine-a mutants, the initial proliferation in each of these tissues is significantly diminished or absent. Regeneration of the caudal fin and extraocular muscle is delayed; regeneration of the retina is nearly completely absent. These data demonstrate that Midkine-a is universally required in the signaling pathways that convert tissue injury into the initial burst of cell proliferation. Further, these data highlight differences in the molecular mechanisms that regulate epimorphic regeneration in zebrafish.

## Introduction

Epimorphic regeneration is the process of replacing ablated cells and tissues, which are then functionally integrated into the mature organ. This process is characterized by the formation of a proliferative blastema at the wound plane, which is capable of fully reconstructing the missing tissues[1]. The regenerative blastema can originate from resident, tissue-specific stem cells or extant mature cells that are reprogrammed into a dedifferentiated state[2,3]. The abiding scientific interest in epimorphic regeneration is sustained by the striking dichotomy in the regenerative abilities between vertebrates, such as amphibians and teleost fish, and mammals[4,5]. Further, identifying the molecular mechanisms that govern epimorphic regeneration holds the promise of informing therapeutic approaches for treating injuries in humans.

Zebrafish is an excellent model to study epimorphic regeneration. This teleost fish has the ability to regenerate multiple tissues, including fins, somatic muscle, heart muscle, and the central nervous system[6–8]. Following amputation, the caudal fin regenerates from intra-ray mesenchymal stem and progenitor cells and dedifferentiated osteoblasts[9–12]. Following ablation, muscle regenerates from dedifferentiated myocytes[10,13,14,15].

In contrast to fin and muscle, where injury reprograms extant cells into tissue-specific progenitors[16,17], regeneration in the central nervous system of zebrafish is sustained by radial glia, which also function as intrinsic neuronal stem cells[8,18–20]. In the retina, Müller glia are the intrinsic stem cells[21]. In response to cell death, Müller glia dedifferentiate, enter the cell cycle, and undergo a single asymmetric division to produce rapidly dividing, multipotent progenitors that continue to divide and differentiate into all types of retinal neurons[22,23]. Cell death also accelerates proliferation of rod precursors that are derived from Müller glia and that contribute genesis of rod photoreceptors[24–27].

Midkine is an evolutionarily conserved, heparin binding cytokine growth factor that in vertebrates has multiple functions during development, tissue repair, and disease[28–30]. During embryonic development in mammals, Midkine is highly expressed in proliferative cells, then rapidly downregulated at mid-gestation[31]. In adults, injuries in a variety of tissues strongly induce reexpression of Midkine, suggesting a universal function of Midkine during tissue injury, repair or regeneration[31–34]. *Midkine-a*, one of two paralogous *midkine* genes in zebrafish, is induced during the regeneration of heart[35], fin[36], skeletal muscle[14] and retina[37,38]. During the selective regeneration of photoreceptors, Midkine-a is expressed by Müller glia, and it is required by these cells to progress from G1 to S phases of the cell cycle[39]. The function of Midkine-a in zebrafish during the regeneration of somatic tissues and following other retinal injury paradigms has not been elucidated.

Using a Midkine-loss of function mutant[39], we compared the injury-induced proliferation and regeneration of three different tissues: caudal fin, extraocular muscle and retina. In the absence of Midkine-a, the initial proliferative response following injury to the caudal fin and extraocular muscle is significantly diminished. In contrast, following ablation of retinal neurons, proliferation is nearly absent, resulting in the failure of regeneration. These results demonstrate that Midkine-a governs the proliferative response in all forms of epimorphic regeneration and highlights differences in the cellular requirements for this injury-induced molecule.

## Materials and Methods

### Animals

Fish were maintained at 28^0^ C on a 14/10 hours light/ dark cycle, using standard husbandry procedures. AB wildtype (*Danio rerio*), *mdka^mi5001^*, *Tg*(*α-actin:mCherry*) and *mdka^mi5001;^ Tg*(*α-actin:mCherry*) of either sex were used at 6 to 12 months of age. All animal procedures were approved by the Institutional Animal Care and Use Committee (IACUC) at the University of Michigan and Wayne State University.

### Lesion Paradigms and labeling of proliferative cells

#### Fin amputation

Fish were anesthetized, and the distal portion of the caudal fins were amputated proximal to the first lepidotrichia branching point. Following amputation, fish were revived and returned to the housing system. All fin amputations were perpendicular to the anterior/posterior plane to avoid uneven fin outgrowth from the dorsal or ventral halves of the fin. Experimental fins were imaged prior to amputation, and at 4, 6, 12, 19, and 32 days post amputation (dpa). For the proliferation assay, fish were immersed in a 5 mM BrdU solution from 3 to 4 dpa as previously described[40]

#### Partial resection of extraocular muscle

Adult fish were anesthetized, and approximately 50% of one lateral rectus muscle was surgically excised[13]. To visualize extraocular muscle *in situ*, lesions were performed in Tg(*α-actin:mCherry*) and *mdka^mi5001;^Tg*(*α-actin:mCherry*) lines. To label proliferating cells, fish were intraperitoneally injected with 20 μL of 10 mM EdU diluted in PBS at 20 hours following injury and sacrificed at 24 hours post injury.

#### Retinal lesions

Retinal neurons were ablated using either a mechanical stab[41] or an intraocular injection of the Na/K-ATPase inhibitor, ouabain. Briefly, mechanical injuries were performed by inserting a 30-gage needle into the dorsal aspect of the eyes through the sclera. Low doses of ouabain were used to selectively kill inner retinal neurons. For this, 0.5 μL of a 5 μM ouabain solution, diluted in PBS, was injected into the intravitreal space [22,42,43]. The contralateral, control eye received an injection of PBS. To assay for cell proliferation, fish were housed for 24 hrs in 5 mM BrdU between days 3 and 5 following the ouabain injection.

### Tissue processing

Fins were harvested at 4 days post amputation (dpa) and were fixed in a 9:1 absolute ethanol and 37% formaldehyde solution. The tissues were infiltrated and frozen in tissue freezing medium (TFM, General Data Company, Cincinnati, OH;[44]). Radial cryosections were cut at 15 μm and mounted on glass slides.

Extraocular muscles were fixed *in situ* in buffered 4% paraformaldehyde (PFA) and the cranium was decalcified using Morse’s solution[13]. 12 μm cryosections through the muscles, eyes, and skull were mounted on glass slides.

Eyes were fixed in either 9:1 ethanolic formaldehyde (100% ethanol: 37% formaldehyde) or buffered 4% PFA overnight at 4C^0^[38,45,46]. All eyes were cryopreserved, embedded in freezing medium, and retinal sections were mounted on glass slides.

### Immunohistochemistry and EdU detection

To visualize BrdU or PCNA, antigen retrieval was first performed using sodium citrate buffer (10 mM sodium citrate, 0.05% Tween-20, pH 6.0) as previously described[38,40,45]. Sections were then incubated in the primary antibody overnight at 4^0^C. After washing with PBST, sections were incubated in secondary antibodies for 1 hour at room temperature. Nuclei were stained with either DAPI or TO-PRO-3. Fluorescence images were captured using a Leica TCS SP5 or SP8 confocal microscope (Leica, Werzler, Germany). To visualize EdU-labelled cells, a Click-iT Assay kit (Thermo Fisher Scientific) was used.

### Cell counts and outgrowth measurement

For fins, BrdU-labeled cells within the distal-most 400 microns of the blastemal compartment were counted. The counts in 3 different blastemas from each fish were averaged (n=5 wildtype and 5 mutants). For lateral rectus muscles, EdU- and DAPI-labeled nuclei were counted and averaged in 4 to 6 non adjacent sections (n=4 wildtype and 4 mutants). In stab lesioned retina, PCNA-positive cells in three non-adjacent sections were counted and averaged in each retina (n=3 wildtype and 3 mutants). In ouabain injected retinas, BrdU-labeled cells were counted in the central dorsal retina in 3 non-adjacent sections per retina (n= 9 wildtype and 9 mutant) and the values were averaged. For outgrowth of the fin, each individual fin was imaged throughout regeneration at multiple time points (pre-amputation, 4, 6, 12, 19, and 32 dpa). Area of fin was calculated using ImageJ software (https://imagej.nih.gov/ij/). Percentage of the outgrowth was obtained from dividing the post-amputation area by the pre-amputation area and multiplied the result by 100.

### qPCR

RNA was prepared from muscle tissue using TRIzol (Invitrogen, Carlsbad, CA), following the manufacturer’s protocol. Each samples of control and injured muscles were pooled from a total of 10 wildtype and 10 *mdka^mi5001^* fish. Following DNase treatment, RNA was quantified using a Nanodrop spectrophotometer (Thermo Fisher Scientific, Waltham, MA). RNA quality was assessed using a 2100 Bioanalyzer (Agilent Technologies, Santa Clara, CA). RNA samples were reverse transcribed using an iScript reverse transcription kit (Bio-Rad Laboratories, Hercules, CA).

Gene expression was measured using CFX96 Real-Time PCR detection system (Bio-Rad Laboratories, Hercules, CA). Diluted cDNA, SsoFast EvaGreen supermix (Bio-Rad Laboratories), and specific primers were used in 20 μl. Reactions were performed in triplicate.

The specificity of the PCR products was verified by analysis of melting curves and by electrophoresis. Gene expression from pools of 3 wildtype and 3 mutants was calculated by the ΔΔ C(t) method[47] using 18S rRNA[48] as the reference gene.

### Statistical analysis

Statistical significance within data sets was analyzed using either the Student *t*-test or one-way analysis of variance (ANOVA) followed by Newman-Keuls multiple comparisons test. All statistical tests were performed using the Prism 6.03 software for Mac OS X (GraphPad Software, Inc. La Jolla, CA). A p-value ≤ 0.05 was considered statistically significant.

## Results

### After an initial delay, Midkine-a mutants completely regenerate fins

Zebrafish regenerate caudal fins from the activation of intra-ray mesenchymal stem and progenitor cells and dedifferentiated osteoblasts[9–12]. RNAseq and gene expression analyses identified *midkine-a* as a gene that is strongly upregulated following fin amputation[36]. To determine the function of Midkine-a during fin regeneration, fins from wildtype and *mdka^mi5001^* were amputated and proliferation was evaluated in each using immunostaining for proliferating cell nuclear antigen (PCNA) and BrdU incorporation. In wildtype animals, the formation and outgrowth of the blastema was observed by 4 dpa (Fig. 1A). Compared with wildtypes, the initial outgrowth of fins in the *mdka^mi5001^* was reduced in size (Fig. 1A-D), and the blastemal compartment contained significantly fewer PCNA- (Fig. 1C-F) and BrdU-positive cells (Fig. 1G-I).

**Figure 1.**
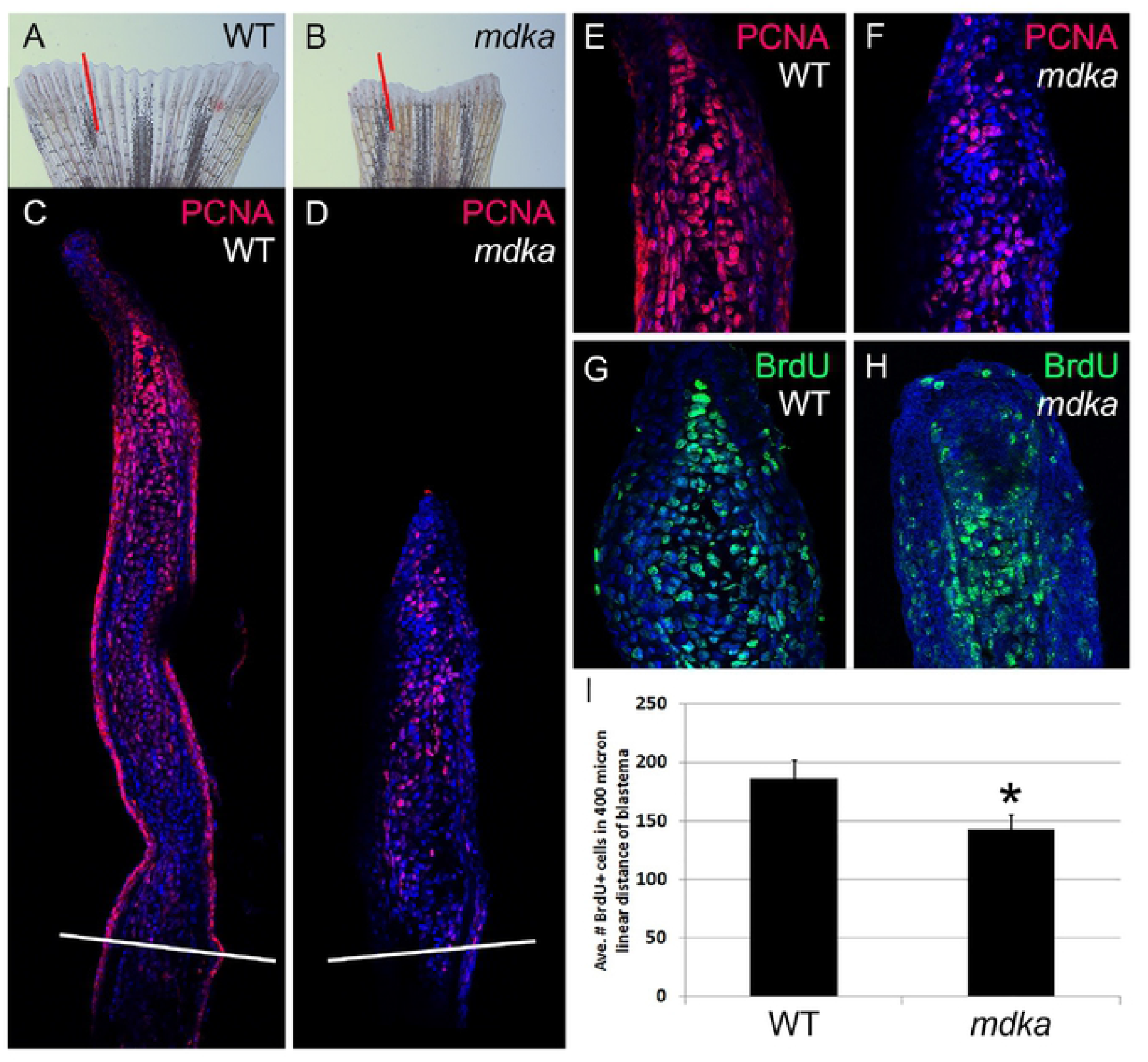
Initial proliferation is diminished in *mdka^mi5001^* following fin amputation. **(A-D)** Stereo and confocal microscope images of regenerating fins in wildtype (A,C) and *mdka^mi5001^* (B,D), immunolabeled with PCNA (magenta). Red lines (A,B) indicate the plane of cross section shown in panels C and D. White lines across the fins show the levels of the amputation plane (C,D). **(E-H)** Confocal images of the blastemal compartments of fins from wildtype (E,G) and *mdka^mi5001^* (F,H), immunolabeled with PCNA (E,F) or BrdU (G,H) incorporated between 3-4 dpa (see Materials and Methods). **(I)** Graph illustrating the average number of BrdU-positive cells within a 400 μm linear distance. Student’s t-test, p=0.0326, n=5 wildtype and 5 mutants.

To determine if the diminished initial proliferation in the *mdka^mi5001^* led to persistently altered regeneration, the area of the regenerated tissue was measured over time. In wildtype animals, regeneration of the caudal fin was completed by 32 dpa (Fig. 2A). Whereas outgrowth of the regenerated fin in *mdka^mi5001^* showed a significant reduction in area at 4 dpa, there was no statistically-significant difference in the area of the regenerated fins between wildtype and *mdka^mi5001^* at the subsequent the time points (Fig. 2A-C). Although the areas of regenerated fins were statistically indistinguishable, regenerated fins in mutants frequently displayed abnormal shapes and mis-patterned melanophore pigmentation (Fig. 2A). These results indicate that Midkine-a is required initially to amplify proliferation among the population of cells that create the blastema, however, once the blastema is formed, Midkine-a is not required, either to sustain proliferation or to complete regeneration. Further, we infer that the structural defects in regenerated fins in *mdka^mi5001^* are a consequence of the initially malformed blastema that is then propagated throughout the period of regeneration.

**Figure 2.**
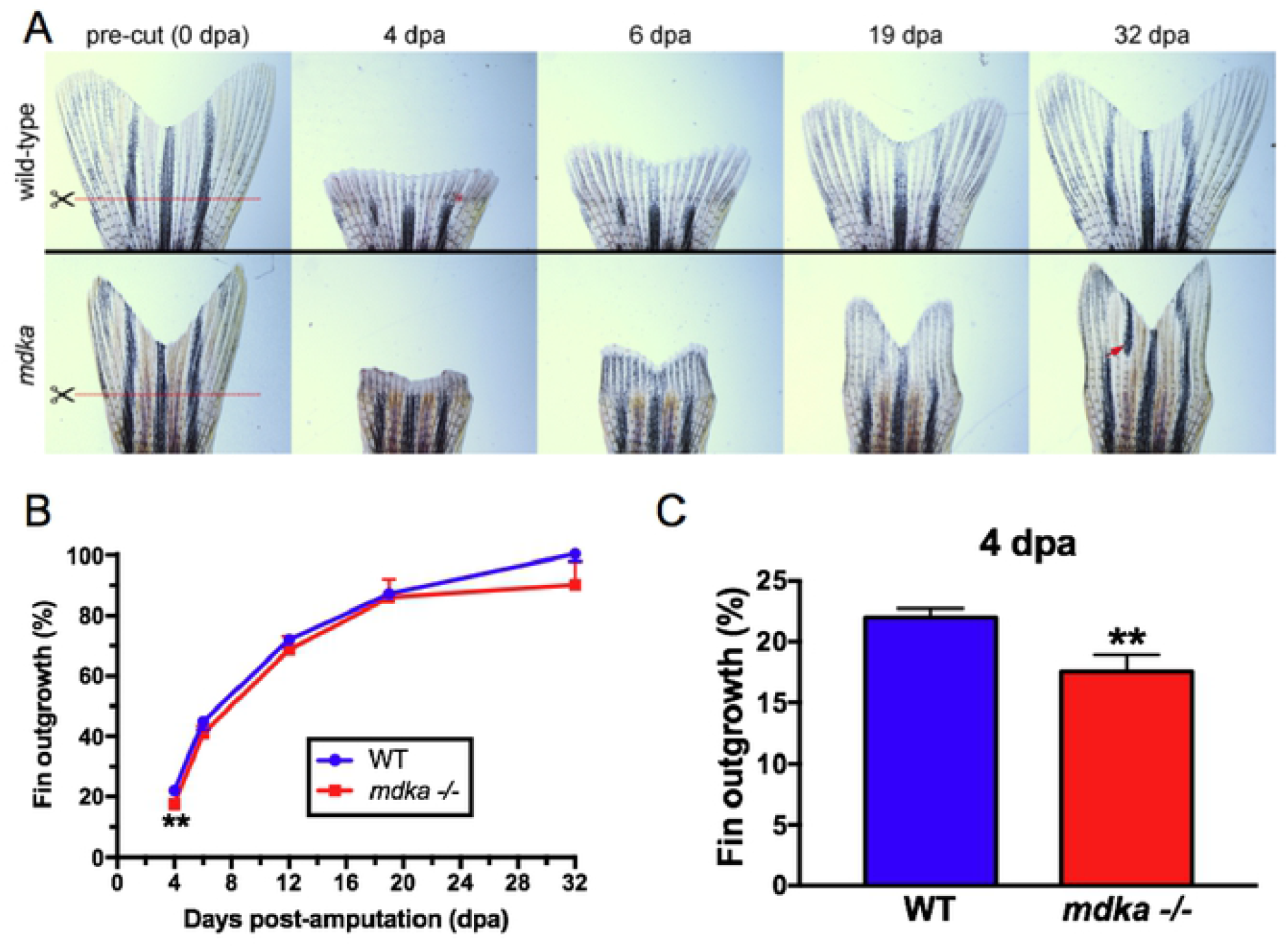
Initial fin regeneration is delayed but recovers in *mdka^mi5001^*. **(A)** Fins from wildtype (top) and *mdka^mi5001^* (bottom) at 0, 4, 6, 19, and 32 dpa. Red lines indicate the level of the amputation plane. In wildtype animals, the structure of the bony fin rays and pigmentation recover completely. In the *mdka^mi5001^*, regenerated fins display abnormal shape and mis-patterned melanophore pigmentation (red arrow). **(B)** Graph illustrating the average percent outgrowth of wildtype and *mdka^mi5001^* through 32 dpa. Wildtype fins recover to 100% of pre-amputation levels by 32 dpa, and mutant animals recover to 90%. **(C)** Graph illustrating the percent fin outgrowth at 4 dpa in wildtype and *mdka^mi5001^*. Student’s t-test, p=0.0105. n = 9 wildtype and 8 mutants.

### In Midkine-a mutants, proliferation and regeneration of extraocular muscle is impaired

In zebrafish, regeneration of extraocular muscles utilizes the dedifferentiation and proliferation of extant myocytes, which then differentiate into muscle cells[10,13,14,48–50]. An RNAseq screen previously showed the upregulation of *midkine-a* during regeneration of extraocular muscle[14]. This result was validated, and it was determined that at 48 hours post lesion (hpl) there is a significant induction of *midkine-a* in the residual stump of muscle (Fig. 3A). Compared with wildtype animals, at 24 hpl the proportion of the EdU-labeled cells in the *mdka^mi5001^* is significantly reduced (Fig. 3B,C), consistent with the early requirement of Midkine-a during regeneration of the fin (see above). Measurements of the muscle growth revealed a significant initial delay in regeneration in the *mdka^mi5001^* (Fig. 3D,E). These results indicate that, similar to regeneration of the caudal fin, Midkine-a is required for the initial proliferation in muscle induced by an injury.

**Figure 3.**
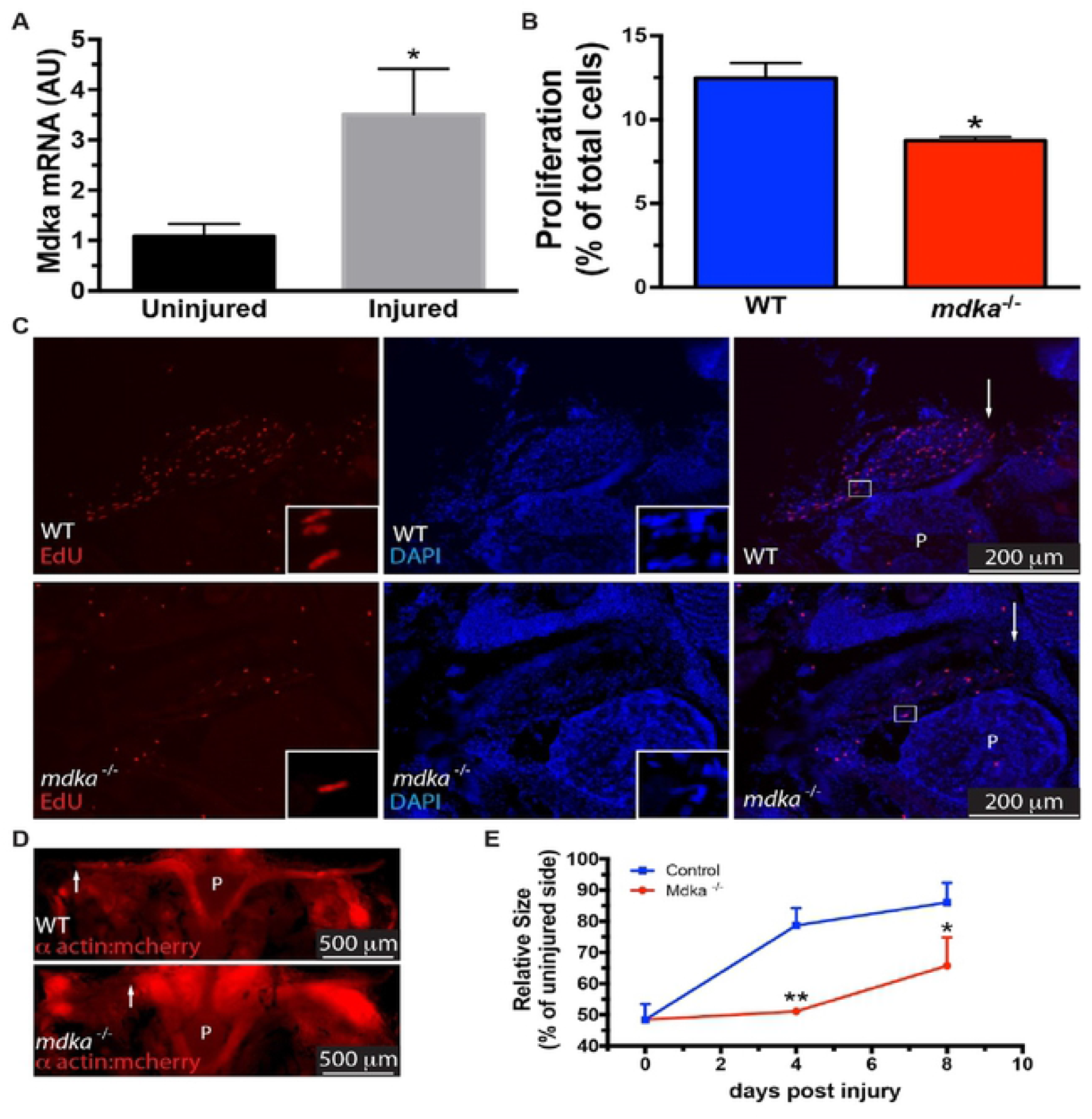
Regeneration of the extraocular muscle is delayed in *mdka^mi5001^*. **(A)** qPCR for *midkine-a* in wildtype zebrafish at 48 hours post lesion. Student’s t-test, p*<0.05, n = 3 wildtype and 3 mutants (10 fish per sample). **(B)** Cell counts of EdU labeled cells in muscles at 24 hours post lesion from wildtype (n=4) and *mdka^mi5001^* (n=4). Student’s t-test, p*<0.05. **(C)** EdU-labeled cells in muscles from wildtype (top) and *mdka^mi5001^* (bottom) at 24 hours post lesion. Arrows indicate the growing tip of the regenerating muscle. P indicates the pituitary body at the midline. **(D)** Images of regenerating extraocular muscles in the Tg(*α-actin:mCherry*) and *mdka^mi5001;^Tg*(*α-actin:mCherry*) lines at 4 days post lesion. **(E)** Outgrowth of regenerating muscles at post-injury time points in wildtype and *mdka^mi5001^*. One-way ANOVA followed by Newman-Keuls multiple comparisons test, p*<0.05, p**<0.01, n = 4 wildtype and 4 mutants.

### Proliferation of Müller glia is diminished in Midkine-a mutants

A recent study demonstrated that, following selective ablation of photoreceptors, Midkine-a is required for Müller glia to progress through the cell cycle[39]. We first confirmed this previous finding using a stab wound lesion, which locally ablates all retinal neurons[51]. In wildtype animals, the proliferation of Müller glia and Müller glia-derived progenitors, which originate in the inner nuclear layer, increases at 48-72 hpl (Fig. 4A,B). However, in the Midkine-a mutants at 48 and 72 hpl, the number of PCNA-positive cells was significantly less (Fig. 4C,D,E,F,G).

**Fig. 4.**
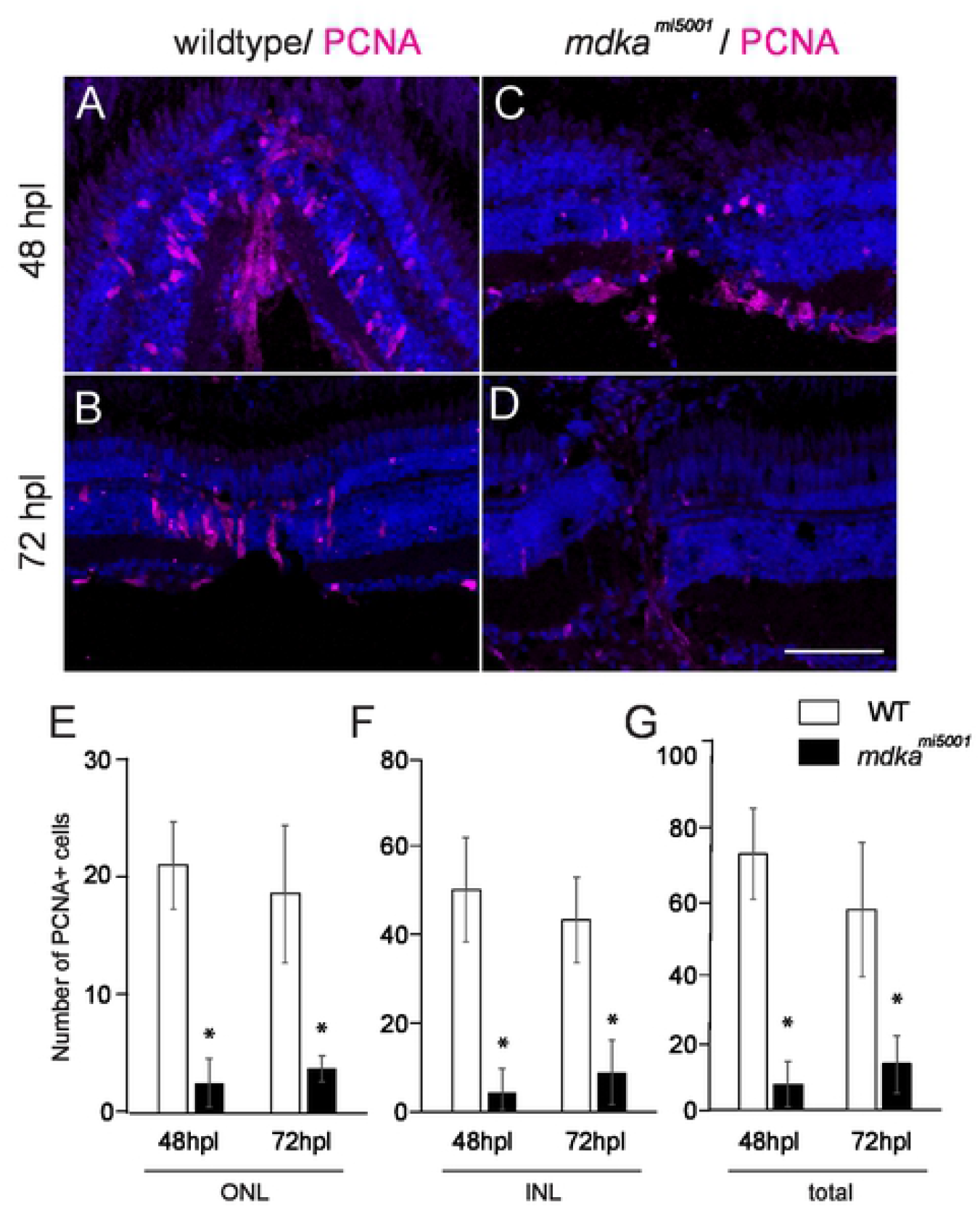
Proliferation of Müller glia is compromised following a stab wound in *mdka^mi5001^*. **(A-D)** Immunocytochemistry for proliferative marker, PCNA, in wildtype (A,B) and *mdka^mi5001^* (C,D) at post-lesion time points. **(E-G)** Graphs illustrating the number of PCNA-positive cells in the outer nuclear layer (E; 48 hpl, wildtype: 20.9 ± 3.7 cells, *mdka^mi5001^*: 2.6 ± 2.1 cells, p=0.0016; 72 hpl, wildtype: 18.5 ± 5.8 cells, *mdka^mi5001^*: 3.8 ± 1.1 cells, p=0.0126), inner nuclear layer (F; 48 hpl, wildtype: 50.7 ± 11.9 cells, *mdka^mi5001^*: 4.4 ± 5.3 cells, p=0.0036; 72 hpl, wildtype: 43.7 ± 9.7 cells, *mdka^mi5001^*: 8.9 ± 7.3 cells, p=0.0077), and total retinal layer (G; 48 hpl, wildtype: 72.1 ± 12.9 cells, *mdka^mi5001^*: 7.1 ± 6.3 cells, p=0.0014; 72 hpl, wildtype: 56.4 ± 19.1 cells, *mdka^mi5001^*: 12.8 ± 8.0 cells, p=0.0212). Scale bar equals 60 μm. Student’s t-test, p*<0.05, n=3 wildtype and 3 mutants.

We also used intraocular injection of low-dose ouabain, to selectively ablate inner retinal neurons[42]. These studies were performed to evaluate this lesion paradigm in the Midkine-a mutants and to determine if rod progenitors, which are spared by ouabain injections[52] and are proposed to be lineage-restricted [39], are capable of regenerating inner retinal neurons. The cell-type specific marker, HuC/D[22,42,53] and PCNA labeling were used to evaluate the ouabain lesions and initial proliferative response, respectively. At 72 hours post injection (hpi), compared with controls, ouabain dramatically reduced the number of HuC/D-positive cells, in both wildtype and *mdka^mi5001^* retinas, demonstrating that low-dose ouabain injections successfully ablate inner retinal neurons (Fig. 5A-C). Consistent with previous reports[43], photoreceptors in the outer retina are largely spared as evident by the normal lamination and cellular density within the outer nuclear layer (Fig. 5A,B). In wildtype retinas at 72 hpi, elongated, PCNA-positive nuclei of Müller glia and Müller glia-derived progenitors are observed in the inner nuclear layer (Fig. 5D). In contrast, at 72 hpi in the *mdka^mi5001^* retinas, very few PCNA positive cells are present in the inner retina (Fig. 5E,F). The PCNA-positive cells present in the mutant retinas are likely either endothelial cells or activated microglia, which are also PCNA-positive in wildtype retinas[22,54]. These results indicate that in the *mdka^mi5001^*, Müller glia fail to proliferate following ablation of inner retinal neurons, and that the extent of cell death or types of retinal neurons ablated does not alter the requirement of Midkine-a for Müller glia to progress through the cell cycle.

**Fig. 5.**
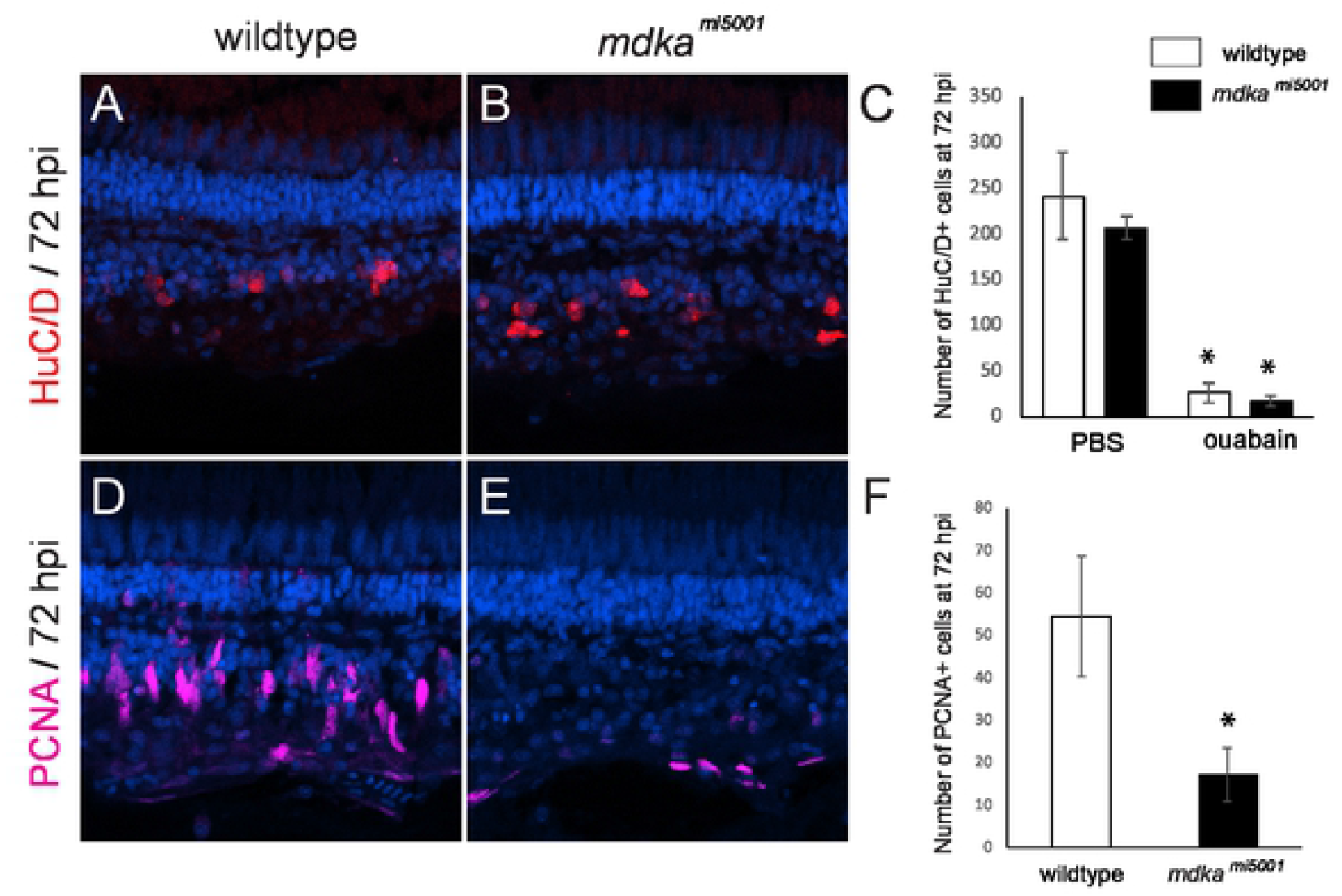
Absence of proliferation in *mdka^mi5001^* following the death of inner retinal neurons. **(A, B)** Immunocytochemistry for the cell marker, HuC/D, in wildtype (A) and *mdka^mi5001^* (B) at 72 hpi. **(C)** The number of HuC/D-positive cells following the injection of PBS (wildtype: 241.3 ± 28.2 cells, *mdka^mi50011^*: 206.0 ± 12.4 cells) and ouabain (wildtype: 25.8 ± 10.5 cells, *mdka^mi5001^*: 17.2 ± 7.2 cells, p=0.137) retinas. **(D, E)** Immunocytochemistry for proliferative marker, PCNA in wildtype (D) and *mdka^mi5001^* (E) at 72 hpi. **(F)** The number of PCNA-positive cells per 500 μm in wildtype (54.4 ± 14.2 cells) and *mdka^mi5001^* (17.1 ± 6.4 cells. p=0.00246) at 72 hpi. Student’s t-test, p*<0.05, n=9 wildtype and 9 mutants.

In addition, animals were exposed to BrdU between 3 and 5 days post injection (dpi) and regenerated neurons were counted at 28 dpi. In wildtype retinas, BrdU-labeled neurons were scattered across all retinal layers, indicating regeneration of inner retinal neurons and a few photoreceptor cells (Fig. 6A). In contrast, there were significantly less BrdU-labeled cells in the *mdka^mi5001^* retinas (Fig. 6B-C). Further, as a consequence of the diminished initial proliferation, at 28 dpi in the *mdka^mi5001^* there was a complete absence of the inner retinal layers (Fig. 6D-F). This striking result also demonstrates that rod progenitors are incapable of regenerating inner retinal neurons (see Nagashima et al., 2020), demonstrating that the fate of rod progenitors is restricted, and these cells are capable only of generating rod photoreceptors.

**Fig. 6.**
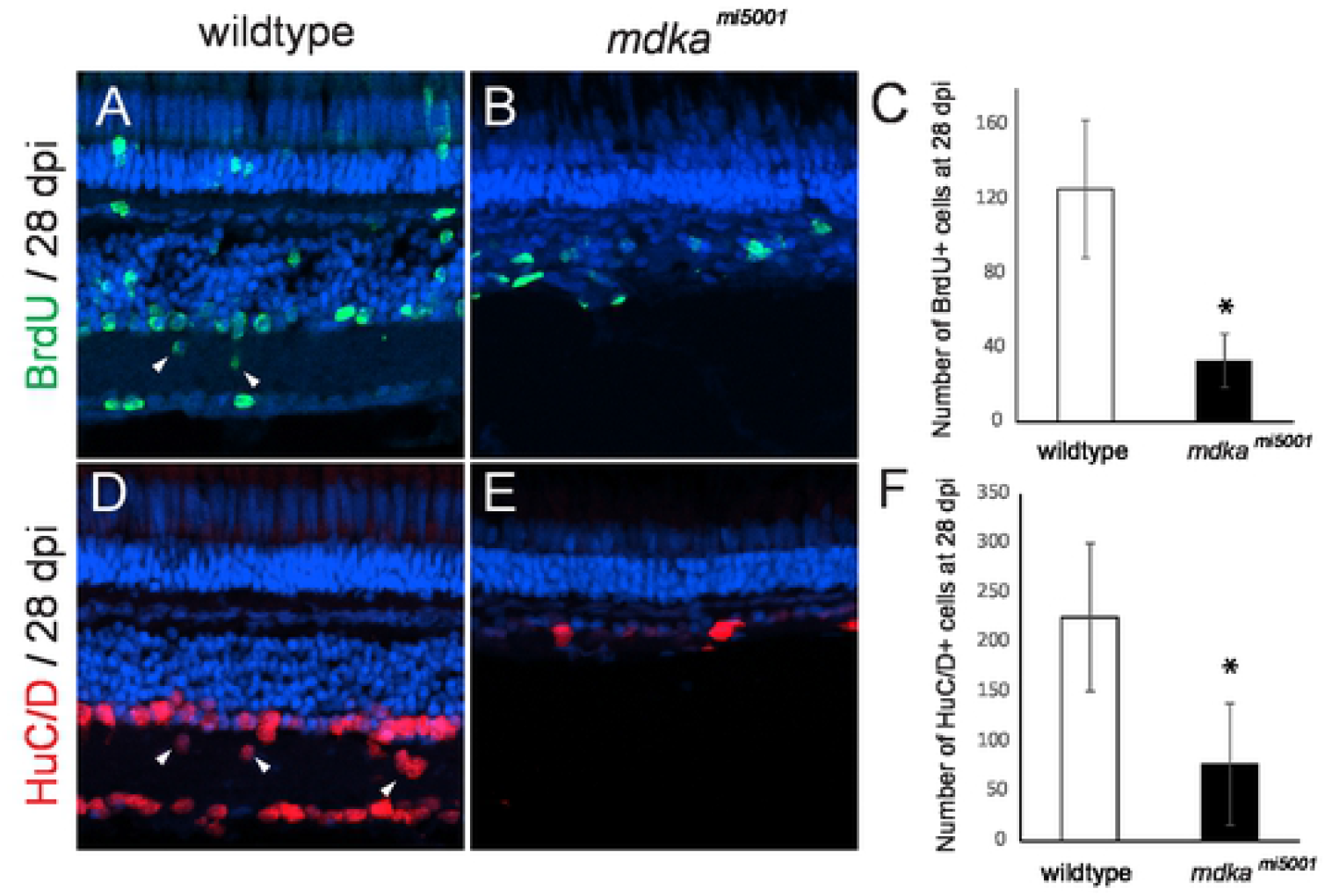
Absence of retinal regeneration in *mdka^mi5001^* at 28 dpl following the death of inner retinal neurons. **(A, B)** BrdU-labeled regenerated retinal neurons in wildtype (A) and *mdka^mi5001^* (B) retinas at 28 dpi. **(C)** The number of BrdU-positive cells wildtype (125.2 ± 37.1 cells) and *mdka^mi5001^* (33.2 ± 14.4 cells, p=0.000341) retinas at 28 dpi. **(D,E)** Regenerated inner neurons labeled with HuC/D in wildtype (D) and *mdka^mi5001^* (E) at 28 dpi. Arrowheads indicate regenerated inner neurons are displaced in the inner plexiform layer. **(F)** The number of HuC/D cells in wildtype (224.8 ± 74.6 cells) and *mdka^mi5001^* (76.8 ± 61.4 cells, p=0.00021) at 28 dpi. Student’s t-test, p*<0.05, n=9 wildtype and 9 mutants.

## Discussion

During epimorphic regeneration, mature, differentiated cells reprogram into stem and progenitor states[55,56]. Proliferation is a fundamental mechanism that serves to amplify these stem and progenitor cells in order to replace lost or damaged tissues. Our results demonstrate that Midkine-a, a soluble cytokine growth factor induced by tissue damage, governs elements of this proliferation during epimorphic regeneration (Fig. 7).

**Fig. 7.**
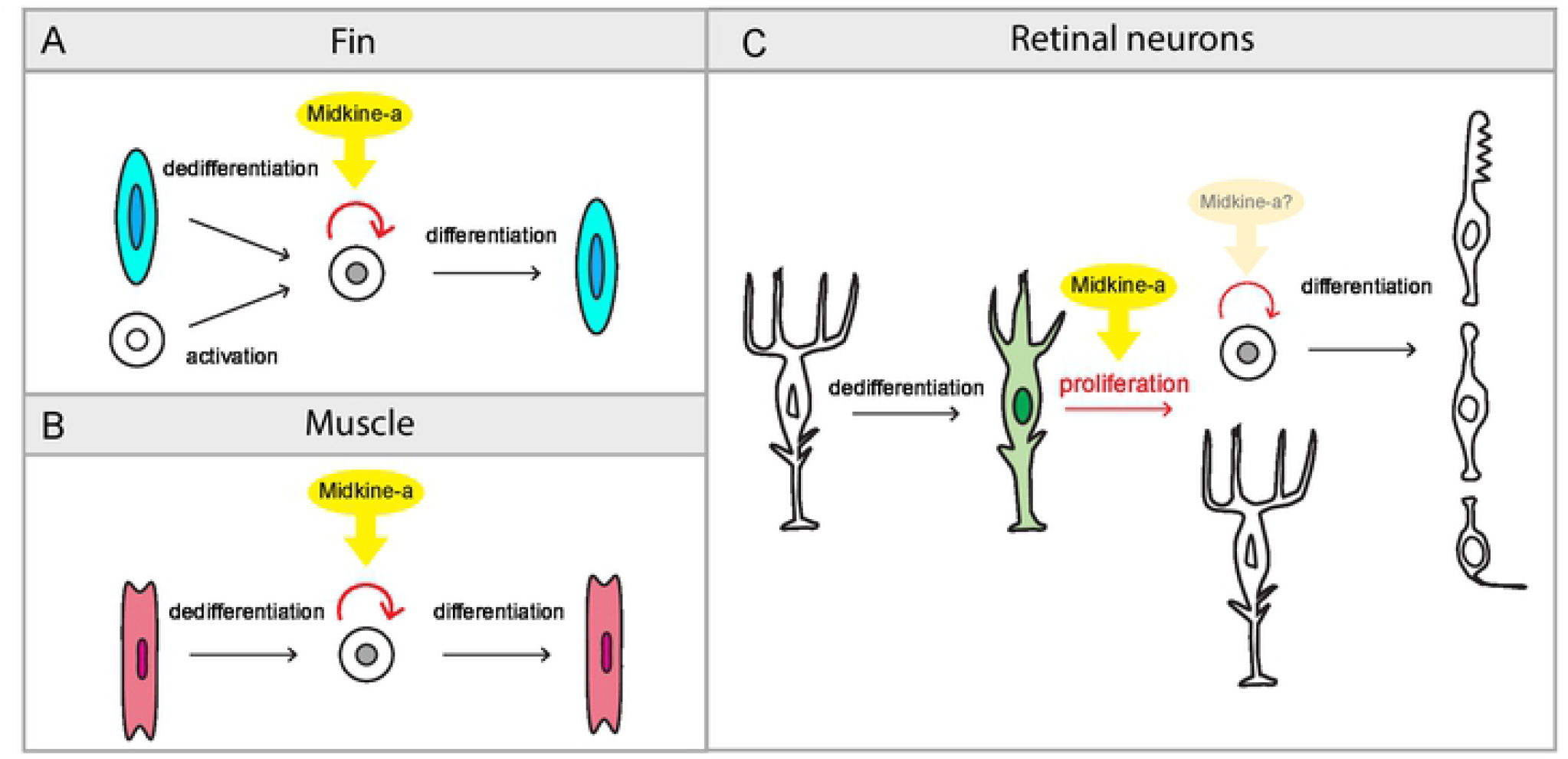
Summary diagram: The function of Midkine-a during epimorphic regeneration. **(A,B)** Following fin amputation, Midkine-a regulates proliferation of precursors produced from dedifferentiation of osteoblasts and activation of mesenchymal stem and progenitors (A). Following extraocular muscle excision, Midkine-a regulates proliferation of precursor produced from dedifferentiation of myocyte (B). During fin and muscle regeneration, loss of Midkine-a results in diminished initial burst of proliferation. **(C)** During regeneration of retinal neurons, Midkine-a governs proliferation of dedifferentiated Müller glia. In the absence of Midkine-a, retinal neurons fail to regenerate.

Regeneration is a multi-step process involving dedifferentiation, proliferation, and differentiation. Numerous factors and signaling pathways regulate different components of this biological process[57,58]. During the regeneration of both fin and extraocular muscle, Fgf signaling governs the formation of the blastema and the proliferation of dedifferentiated cells[50,59,60]. Similarly, during regeneration of the fin Igf signaling is required for the proliferation of blastema cells. In contrast, during the regeneration of muscle Igf is involved late, during the differentiation of newly generated myoblasts[49,61]. Our results demonstrate that the cytokine growth factor, Midkine-a, is also involved in tissue regeneration, possibly as a generic wound signal to regulate initial burst of proliferation during the regeneration of both fin and muscle (Fig. 7A,B and[62]). Although the cell types in fin and muscle that express Midkine-a are not yet established, we infer that in these tissues Midkine-a functions as a paracrine and/or autocrine regulator of proliferation that acts on activated, dedifferentiated cells that give rise to the regeneration blastema.

There are at least two mechanisms that can explain the function of Midkine-a during the regeneration of fin, muscle, and retinal neurons. First, Midkine-a may be required for a subset of reprogrammed cells to enter into, or progress through the cell cycle. Cellular reprogramming occurs in all the tissues examined, and entering and/or progressing through the cell cycle is a universal requirement among reprogrammed cells. During regeneration of fin, proliferative cells in the blastemal are derived from multiple sources of reprogrammed cells, whereas during muscle regeneration myocytes are solely responsible to form the blastema. The initial delay of the burst of proliferation may reflect incomplete contribution among cells capable of forming the blastema. This scenario is consistent with that following light lesions, very few Müller glia are able to progress through the cell cycle[39]. Second, Midkine-a may govern cell cycle kinetics among progenitors derived from reprogrammed and/or stem-like cells. Midkine-a may be required for the amplification of this population of cells, perhaps serving to match the initial size of the blastema to the spatial extent of the injury. Utilizing murine embryonic stem cells[63], human cancer-derived cell lines[64], and the embryonic retina in zebrafish[65], it has been established that Midkine can govern cell cycle kinetics. Additional research will be required to determine which of these alternative mechanisms is correct.

In the intact teleost retina, rod precursors give rise to rod photoreceptors[24–26]. Our data are consistent with a previous report that in the Midkine-a mutants following selective ablation of photoreceptors, the regeneration of cone photoreceptors is permanently compromised, whereas rod photoreceptors are replenished (38). The failure to regenerate inner retinal neurons in the Midkine-a mutants indicates that the fate of rod precursors is restricted to the rod photoreceptor lineage. Given that rod precursors originate from Müller glia[18], this result suggests that following their migration to the outer retina a lineage restriction is imposed on these cells by the local environment within the outer nuclear layer. Our data also demonstrate that Midkine-a-mediated proliferation of Müller glia and generation of Müller glia-derived progenitors are required for regeneration of retinal neurons. This conclusion leads to the question: Does Midkine-a regulate proliferation of Müller glia-derived progenitors (Fig. 7C)? The current manuscript does not investigate the question, however, we would predict two possible scenarios. First, in the mutants, proliferation of progenitors is also blocked. This would indicate conserved molecular machinery to regulate cell cycle among the cells that proliferate and Midkine-a plays a major role during the proliferation. Second, in the absence of Midkine-a, proliferation of progenitors is largely intact. This would imply that a different set of factors are required for proliferation of dedifferentiated cells versus progenitors. If the cells are already in the active cell cycle, Midkine-a may play a less role. We favor the second possibility, supporting previous observation that in the absence of Midkine-a, mutant animals progress slowly but normally through development and post-embryonic growth[39]. Additional studies will be needed to further understand cell type specific requirements of Midkine-a during proliferation.

Taken together, our data demonstrate that in zebrafish Midkine-a universally regulates the initial proliferation of reprogrammed cells and/or their progenitors (see also 38), which are activated following tissue damage. Further, our results highlight differences in the molecular mechanisms by which Midkine-a regulates epimorphic regeneration. It remains to be determined if variability in receptors or intracellular signaling pathways is responsible for the differences in the requirement of Midkine-a during the regeneration of different tissues.

## Acknowledgements

This work was supported by grants from the National Institutes of Health (NEI); R01EY07060 (to PFH) and R01EY026551 (to RT). Histology and imaging core resources were supported by vision core grants (P30EY07003 and P30EY04068) and an unrestricted grant from Research to Prevent Blindness to each department. Fish lines and reagents provided by ZIRC were supported by NIH-NCRR Grant P40 RR01. The authors thank Dilip Pawar and Xixia Luo for zebrafish maintenance and technical assistance.

